# NUMT PARSER: automated identification and removal of nuclear mitochondrial pseudogenes (numts) for accurate mitochondrial genome reconstruction in *Panthera*

**DOI:** 10.1101/2022.04.04.487049

**Authors:** Alida de Flamingh, Angel G. Rivera-Colón, Tom P. Gnoske, Julian C. Kerbis Peterhans, Julian Catchen, Ripan S. Malhi, Alfred L. Roca

**Author notes:** Corresponding author/s: Alida de Flamingh & Angel G. Rivera-Colón. Equal contributing first authors (Alida de Flamingh, Angel G. Rivera-Colón).

## Abstract

Nuclear mitochondrial pseudogenes (numts) may hinder the reconstruction of mtDNA genomes and affect the reliability of mtDNA datasets for phylogenetic and population genetic comparisons. Here, we present the program Numt Parser, which allows for the identification of DNA sequences that likely originate from numt pseudogene DNA. Sequencing reads are classified as originating from either numt or true cytoplasmic mitochondrial (cymt) DNA by direct comparison against cymt and numt reference sequences. Classified reads can then be parsed into cymt or numt datasets. We tested this program using whole genome shotgun-sequenced data from two ancient Cape lions (*Panthera leo*), because mtDNA is often the marker of choice for ancient DNA studies and the genus *Panthera* is known to have numt pseudogenes. Numt Parser decreased sequence disagreements that were likely due to numt pseudogene contamination and equalized read coverage across the mitogenome by removing reads that likely originated from numts. We compared the efficacy of Numt Parser to two other bioinformatic approaches that can be used to account for numt contamination. We found that Numt Parser outperformed approaches that rely only on read alignment or Basic Local Alignment Search Tool (BLAST) properties, and was effective at identifying sequences that likely originated from numts while having minimal impacts on the recovery of cymt reads. Numt Parser therefore improves the reconstruction of true mitogenomes, allowing for more accurate and robust biological inferences.

## Introduction

Mitochondrial DNA is often used for phylogenetic studies that investigate matrilineal inheritance patterns (Chaitanya et al., 2014), inter- and intra-specific divergences (Bowers et al., 1994; Cronin et al., 1991; Gill et al., 1993), and for studies that use samples with low DNA copy numbers (Hofreiter, Serre, et al., 2001; Merheb et al., 2019). However, the presence of nuclear mitochondrial (numt) pseudogenes (designated as *Numt* by Lopez et al (Lopez et al., 1994)) may hinder the identification of true cytoplasmic mitochondrial (cymt) DNA sequences and the reliability of mtDNA for phylogenetic and population genetic comparisons (Bensasson et al., 2001; Smart et al., 2019). Numts arise when mitochondrial DNA is incorporated into the nuclear genome during chromosomal double-strand break repair by nonhomologous recombination (Bensasson et al., 2001; Blanchard & Schmidt, 1995, 1996). Organellar DNA fragments are found in the nuclear genomes of many eukaryotes (Bensasson et al., 2001; Gaziev & Shaikhaev, 2010), mostly in non-coding intergenic regions and introns (Bensasson et al., 2001; Gaziev & Shaikhaev, 2010; Smart et al., 2019). Numts may vary in size and sequence depending on the mitochondrial DNA that is integrated into the nuclear genome during double stranded break repair. The nuclear genome integration sites also vary among taxa, with most numts occurring as single copies at dispersed genomic locations (Blanchard & Schmidt, 1995; Zischler, 2000). Some taxa have tandemly repeated numt sequences, e.g., domestic cats (Lopez et al., 1994), while in other taxa numts may be present in telomeric, centromeric and/or other regions of their nuclear genomes (e.g., grasshoppers (Vaughan et al., 1999)).

Primers used to amplify targeted regions of the mitogenome through PCR may also amplify numt pseudogenes, and in some cases may even preferentially amplify numts compared to cymts (Collura & Stewart, 1995). We, and others, e.g., Bensasson et al., 2001, Goios et al., 2008 and Song et al., 2008, use the word *contaminate* to refer to mtDNA datasets where the reads derive from both cymt and numt DNA templates, and where mtDNA datasets are thus contaminated by reads that originate from the nuclear genome. The numt contaminants may show up as “ghost” bands or additional bands in electrophoresis gels post-PCR-amplification (Den Tex et al., 2010), or as sequence ambiguities and nonsense mutations in sequenced DNA, and may result in unexpected phylogenetic placements (Triant & Dewoody, 2009; Zhang & Hewitt, 1996). When amplifying whole genome DNA using high-throughput sequencing technologies that do not target specific genomic regions, numts may cause mitogenome regions with paralogous numt pseudogenes to appear over-represented in a sequence read pool. Numt dataset contamination may therefore be evident when aligning data obtained from whole genome DNA, combining nuclear and mitochondrial reads, to a reference mitogenome. Should DNA amplification be uniform across the genome, then mitogenome regions with corresponding numt pseudogenes will appear to have higher read coverage compared to the mitogenome-wide average, since those regions will be represented in the contaminated dataset by both cymt and numt DNA reads.

Contamination by numts is especially problematic for taxa that have many copies of one or more numt pseudogenes (e.g., cats and other felids have a tandemly repeated 7.9 kilobase pair (kb) numt pseudogene (Lopez et al., 1994)). The higher the numt copy number, or number of tandem repeats, the more the pseudogene will be represented in the shotgun sequence read pool. Taxa that have many numt repeats are therefore especially prone to dataset contamination and incorrect mitogenome characterization.

The unknown presence of numt pseudogenes can lead to incorrect but seemingly convincing phylogenetic results. When numt sequences from one taxon are compared to cymt sequences from other taxa, the resulting phylogenetic inference may not be accurate since the sequences that are being compared are not orthologous. Numt contamination has been a challenge for systematics and phylogenetic inferences of contemporary samples (Allende et al., 2001; Antunes & Ramos, 2005; Pereira & Baker, 2004; Podnar et al., 2007; Sorenson & Quinn, 1998), and has also been especially problematic for studies using ancient DNA (van der Kuyl et al., 1995; Woodward et al., 1994; Zischler, Hoss, et al., 1995) prior to the implementation of ancient DNA validation procedures (Cooper & Poinar, 2000; Gilbert et al., 2005).

Here, we developed and tested a program, Numt Parser, which allows for the identification and filtering of DNA sequences that likely originate from numts in short-read sequencing datasets. Sequencing reads are classified as putatively originating from either cymt or numt DNA by direct comparison against cymt and numt reference sequences. As input, Numt Parser uses sequence alignment map (SAM) files of sequencing reads that have been aligned to cymt and numt reference sequences. Classified reads can then be parsed into separate datasets that contain putative cymt or numt derived reads. We tested Numt Parser using whole genome shotgun-sequenced data from two ancient Cape lions (*Panthera leo*). We focused our analysis on a species within the genus *Panthera* as many taxa within this genus are known to have numt pseudogenes (see section “Numt contamination in the genus *Panthera*”). We also specifically tested the efficacy of Numt Parser on ancient DNA samples that likely contain degraded DNA, as mtDNA is often the marker of choice for ancient DNA studies, and because past studies have found numt pseudogene contamination to be common among ancient DNA samples (Den Tex et al., 2010). We compared Numt Parser to two alternative bioinformatic approaches that may be used to account for numt contamination, including evaluation of standard-practice approaches that rely on alignment-based filtering. We show that Numt Parser outperforms other bioinformatic approaches and is effective at identifying sequences that likely originate from numt pseudogene DNA while having minimal impact on the recovery of cymt reads.

## Methods

### Ancient DNA extraction

We collected and analyzed bone and tooth samples from two Cape lion specimens currently housed at the Field Museum of Natural History (FMNH) in Chicago, USA. Cape lions are thought to have gone extinct in ca. 1850 (Van Bree, 1998). The two skulls studied in this paper were from the colloquially named “Cape Flats” region of South Africa, near the current city of Cape Town on the south-western tip of southern Africa.

In keeping with their current FMNH designation, the Cape lion samples are listed as JCK 10711 and JCK 10712. Samples were collected at the FMNH following a protocol that minimizes contamination by non-target DNA sources. Samples were collected in a dedicated workspace and with equipment that were decontaminated with Takara Bio DNA-off prior to collection of each sample. Disposable personal protective equipment (lab coat, hair net, gloves, cover sleeves) were changed and decontaminated with Takara Bio DNA-off between collection events. The surface of the collection location on each lion specimen skull or tooth was decontaminated with 6% sodium hypochlorite (full strength Clorox bleach) and rinsed with DNA-free ddH_2_O. We collected approximately 0.2 g of bone or tooth powder using a Dremel hand drill that was decontaminated with Takara Bio DNA-Off prior to collection, using a new with a 1mm bit for each collection event.

DNA was extracted in the Malhi ancient DNA laboratory at the Carl. R. Woese Institute for Genomic Biology at the University of Illinois at Urbana-Champaign. The Malhi ancient DNA laboratory is an isolated, air-filtered facility that is dedicated to molecular analysis of ancient and low-template DNA samples. Each tooth and/or bone sample was incubated under rotation in 4 ml of digestion buffer (0.5 M EDTA, 33.3 mg/ml proteinase K, 10% N-lauryl sarcosine) for 12–24 hours at 37°C. Using Amicon K4 centrifugal units, we then concentrated the sample to approximately 250 μl, and used this concentrate as starting template for the DNA extraction using a Qiagen PCR Purification Kit with a final DNA elution volume of 60 μl. Whole genomic libraries were constructed using the NEBNext^®^ Ultra II^TM^ DNA Library Prep kit and NEBNext^®^ Multiplex Oligos (Unique Dual Indexes) for Illumina^®^. The extracted DNA was pre-treated with USER (Uracil-Specific Excision Reagent) enzyme to remove cytosine to uracil nucleotide base changes that are common in ancient DNA (Supplementary file 1 – Step #7A; (Hofreiter, Serre, et al., 2001)).

All samples and negative libraries were pooled, and shotgun sequenced on an Illumina NovaSeq 6000 S1 1×100bp flow cell at the Roy J. Carver Biotechnology Center at the University of Illinois at Urbana-Champaign (see Supplementary file 1 for a step-by-step description of ancient DNA extraction and library preparation protocols).

### Bioinformatic preparation of Numt Parser input files

Samples were de-multiplexed and were merged into a single datafile per lion. Sequence reads were trimmed to have a minimum length of 25 bp using FastP v.0.19.6 (Chen et al., 2018). We assessed whether DNA showed damage patterns that are characteristic of ancient DNA by aligning trimmed reads to the complete lion genome (v. PanLeo1.0, GenBank accession number GCA_008797005.1; Armstrong et al., 2020) using the `mem` module from BWA v. 0.7.15(H. Li & Durbin, 2010) and quantifying damage in mapDamage2 v. 2.0.5 (Jónsson et al., 2013) using a fragment size of 70 bp.

Reads were aligned (Li & Durbin, 2010) to a previously published reference lion mitogenome that showed no evidence of numt contamination, hereafter referred to as the cymt genome (GenBank accession KP202262; see Li et al 2016 for details on validation of mitogenome authenticity). Reads were also aligned to a numt reference sequence (see below) (Figure 1A). Alignments were converted to BAM format and filtered to remove unmapped reads and alignments with a mapping quality less than 30 using SAMtools `view` v. 1.1 (Li et al., 2009). Filtered alignments were then sorted and indexed, with duplicates marked and removed with the Picard Toolkit v. 2.10.1 (“Picard Toolkit.” 2019. Broad Institute). Filtered BAM files were converted to SAM format for compatibility with Numt Parser (Figure 1A).

**Figure 1.**
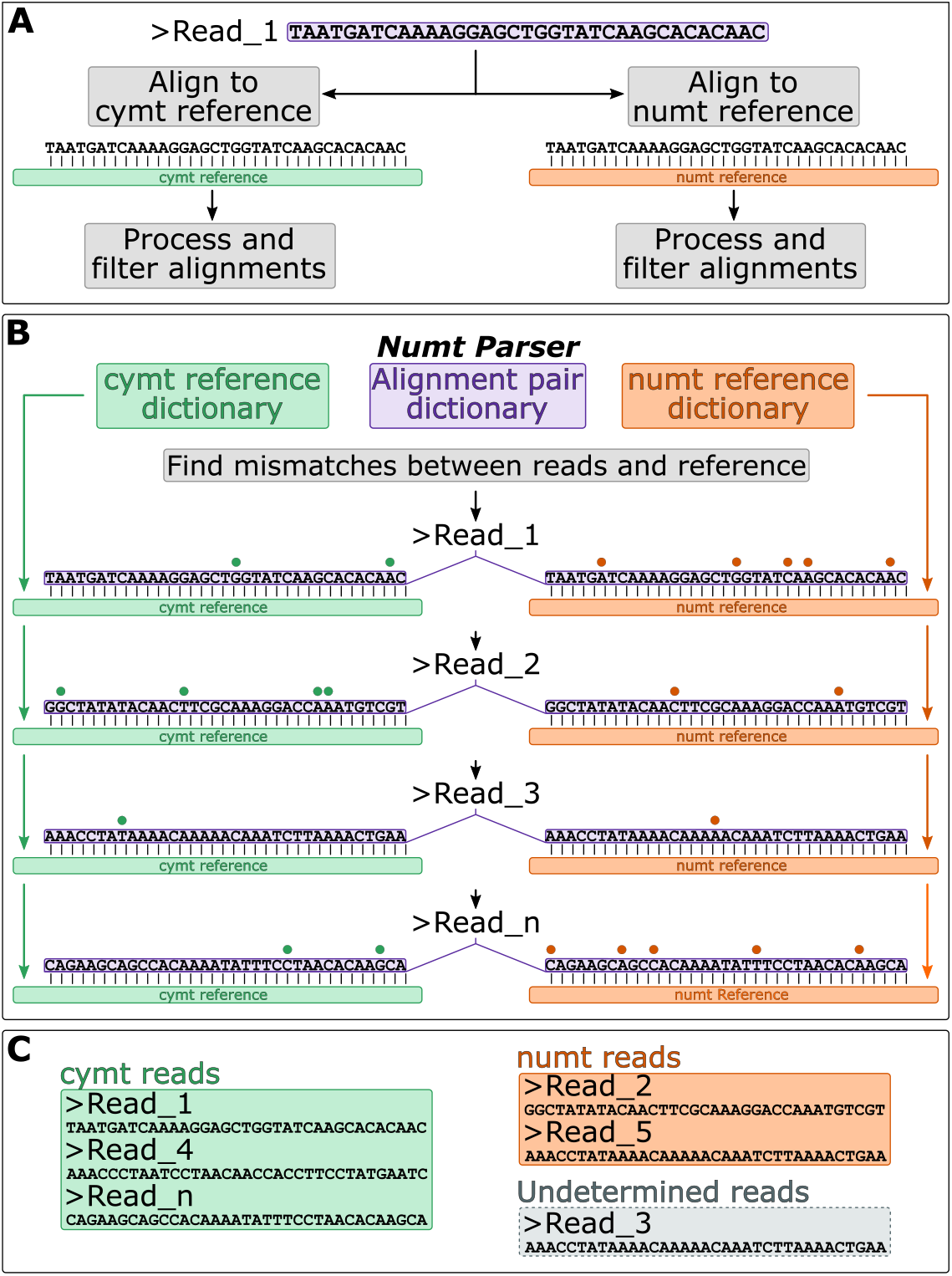
The Numt Parser pipeline for the identification and removal of numts. (A) After demultiplexing and trimming, sequencing reads are independently aligned to a mitogenome reference sequence, and to a reference sequence for numt. The resulting alignments are subsequently filtered and processed by Numt Parser. (B) After alignment, reads are processed by Numt Parser. First, the program reads the reference sequences for both cymt (green) and numt (orange) references and corresponding alignments (purple). The program then iterates over the alignment dictionary, processing each read independently. For each read query, Numt Parser compares the read sequence against the two references and identifies mismatches between reads and each of the references (shown as dots above read alignments). The program outputs a tally of the number of mismatches observed for each read against both the cymt and numt reference sequences. (C) Using the Numt Parser output, the total read pool can then be parsed into datasets containing sequences likely originating from either cymt or numt DNA. Read_n indicates the *n*-th read in the read dataset.

### *Numt contamination in the genus* Panthera

The development and testing of Numt Parser using lion data would be especially pertinent because in taxa within the genus *Panthera*, the presence of numt pseudogenes has interfered with accurate mitogenome reconstruction (Bertola et al., 2016; Curry et al., 2021; Kim et al., 2006; Li et al., 2016). Notably, Li et al. (2016) showed that about half of the published big cat mitogenomes in *Panthera* contain long stretches of disproportionately high sequence divergence consistent with numt contamination. Lopez et al. (1994) showed that some taxa within the family Felidae have a ~7.9 kb numt pseudogene that is tandemly repeated 38-76 times and comprises a macrosatellite with repeats of varying lengths. Using a 12S mtDNA fragment, Lopez et al. (1994) showed that this numt originated before the radiation of modern cat species within the genus *Felis*. Kim et al. (2006) used targeted PCR and DNA cloning to characterize a large numt found in various species of the genus *Panthera*. They identified the cytogenic location of the mtDNA translocation using *in-situ* hybridization and compared the gene sequences of cymt and numt for 5 species of *Panthera.* Their results support two independent translocations of mtDNA into the nuclear genome; one in *Panthera* around 3 million-years ago (MYA) and the other (identified by Lopez et al 1994) into the ancestor of the domestic cat lineage around 1.8 MYA.

### Specifying the numt reference sequence in lion

Li et al. (2016) previously identified nine published felid mitogenomes in GenBank that proved to be composites of both cymt and numt sequences. In the case of the African lion, Li et al. (2016) reported a 7.2 kb region in a published mitogenome sequence (GenBank accession number KF907306) that showed disproportionately high sequence divergence when compared to the lion reference mitogenome (KP202262) and to other lion mitogenome sequences (KC834784 and JQ904290). The patterns of elevated divergence, limited to this 7.2 kb region, indicated that the large numt pseudogene present in *Panthera* had also been mistakenly incorporated into the KF907306 mitogenome sequence. Li et al. (2016) established that the numt pseudogene extends approximately from position 4,250 to 11,450 bp of the lion reference mitogenome (KP202262), corresponding to genes *ND1* partial, *ND2*, *COX1*, *COX2*, *ATP8*, *ATP6*, *COX3*, *ND3*, *ND4L*, *ND4* partial (Figure 2A). Additionally, several of the genes contained in this 7.2 kb region in KF907306 also exhibit an increase in non-synonymous nucleotide and subsequent amino acid changes (Supplementary Figure 1), providing further support for the presence of numt contamination. We therefore used this previously characterized 7.2 kb region in KF907306 described and published by Li et al. (2016) as the numt reference in Numt Parser.

**Figure 2.**
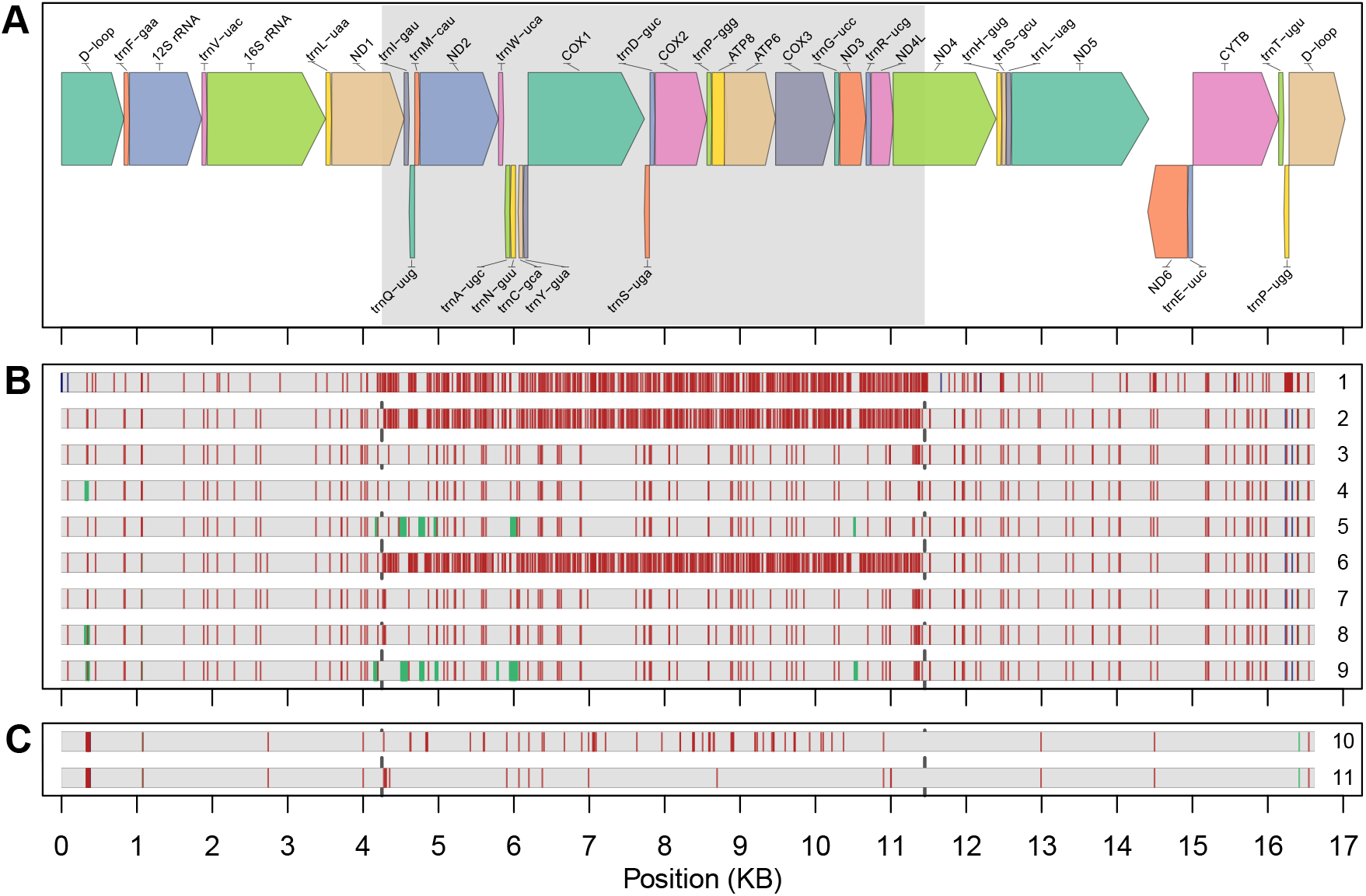
(A) The numt reference for use in Numt Parser was identified by Li et al (2016) and corresponds with positions 4,250 to 11,450 of GenBank accession KP202262 (Li et al., 2016). The region of the numt is homologous to *ND1* (partial), *ND2*, *COX1*, *COX2*, *ATP8*, *ATP6*, *COX3*, *ND3*, *ND4L*, and *ND4* (partial), and is shaded in the figure (the rest of KP202262 corresponded to cymt). GenBank accession KP202262.1, a mitogenome identified as not having numt contamination (Li et al 2016) was used as the cymt reference. (B) Alignments of unfiltered and filtered consensus sequences against the cymt reference mitogenome are shown, with the span of the mitogenome alignment shown as a horizontal grey bar. Vertical red bars represent single-base differences; green bars show disagreements due to uncalled bases (Ns) in the consensus sequence. Blue indicates a deletion in the consensus with respect to the cymt reference. Dark grey dashed lines show the boundaries of the numt pseudogene. Numbers on the right indicate that the following consensus sequence was aligned to the cymt reference: (1) published sequence with known numt incorporation (GenBank accession KF907306; numt reference), (2) unfiltered JCK 10711, (3) Numt Parser-filtered JCK 10711, (4) BLAST-filtered JCK 10711, (5) SAMtools-filtered JCK 10711, (6) unfiltered JCK 10712, (7) Numt Parser-filtered JCK 10712, (8) BLAST-filtered JCK 10712, (9) SAMtools-filtered JCK 10712. In the unfiltered consensus (2 and 6), majority of sequence disagreements are localized within the span of the numt sequence. These disagreements were largely removed by the filtering of the numt-contaminant reads from the consensus. (C) Alignments between the JCK 10711 and JCK 10712 consensus sequences for unfiltered datasets (10) and Numt Parser-filtered datasets (11). Alignment of the unfiltered consensus (10) leads to an increased presence of disagreements between the two consensus sequences for these lions. The disagreements between these consensus sequences overlap with numt pseudogene region consistent with numt contamination. Filtering numt contamination with Numt Parser leads to a reduction in the number of disagreements, but still allows for the retention of sequence differences between the mitogenomes of the two lions (red bars in 11).

### Identifying numt reads with Numt Parser

The Numt Parser analysis (Figure 1B) starts by loading the reference sequences of both the cymt and numt sequences into a reference sequence dictionary that respectively stores their IDs and sequences as key-value pairs. Read alignments in SAM format against both the cymt and numt reference sequences are also loaded and then processed into a separate alignment pair dictionary. Taking each unique read ID as a key, the alignment pair dictionary stores the mapping information (chromosome, base pair, alignment flags, and CIGAR; (Li et al., 2009)) for alignments against both the cymt and numt references. The software then iterates over all reads, comparing them directly to both cymt and numt sequences present in the reference sequence dictionary. Using the corresponding positional information from the alignment, every nucleotide in the read is directly compared against the corresponding site in the reference and a tally is taken of all mismatches between both sequences. Numt Parser accounts for the underlying alignment information and thus handles indels and clipped bases, as determined by the CIGAR string, which are also tallied as mismatches. The number of matches (aligned portion minus tallied mismatches) is then divided by the total length of the aligned portion of the read to obtain a percent identity of each read against each reference. When processing paired-end reads, the comparison between alignment and reference sequences is independently performed for each read in the pair. The Numt Parser output is a table which contains the percent identity, alignment length, and number of mismatches of each unique read ID against both the cymt and numt references. The output also contains the orientation of each read; when the input alignments originate from single-end reads, the read_in_pair column will show a value of 0, since none of the reads are paired. When the alignments originate form paired-end reads, the output table will instead show a value of 1 or 2 in the read_in_pair column to denote the first or second read in the pair, respectively. The output table also contains a classifier that tags each read as putatively originating from cymt DNA (tagged as “cymt”) or from numt DNA (tagged as “numt”) based on the highest percent identity, or tagged as “Undetermined” when reads have equal percent identity to both the cymt mitogenome and numt pseudogene references (Table 1). Provided that the reference sequences and alignments are available, Numt Parser works as a standalone Python program with no requirement of outside software or library dependencies. All required code for running Numt Parser is available at https://github.com/adeflamingh/NuMt_parser.

**Table 1.**
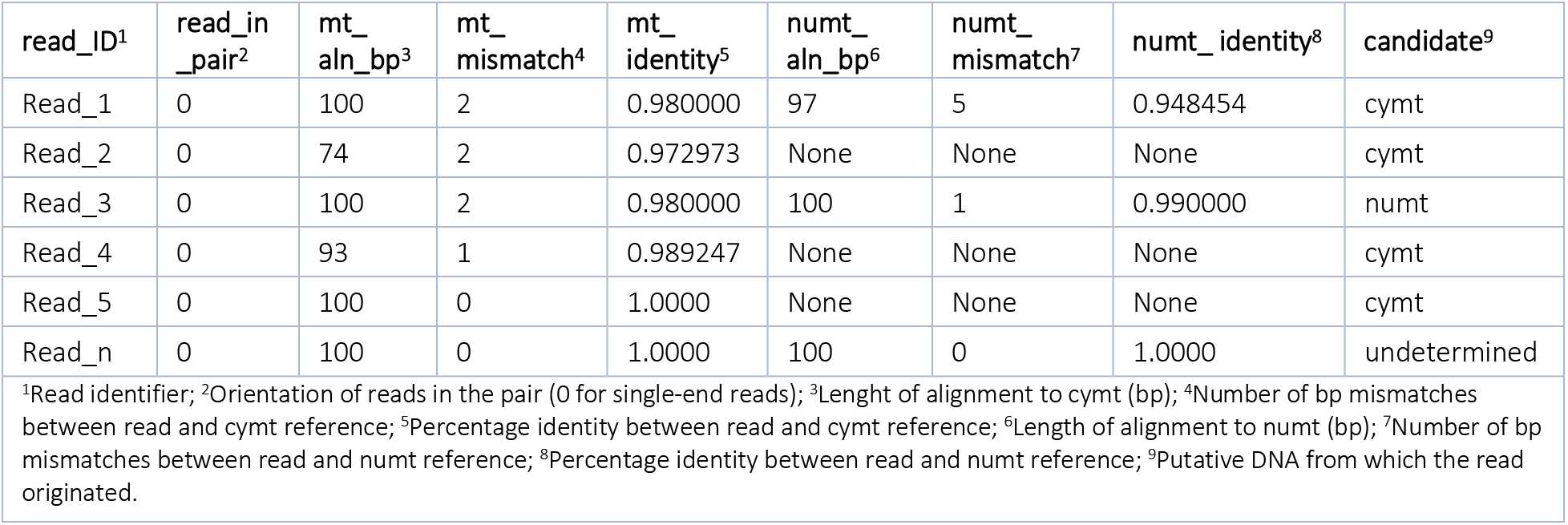
Example table produced by Numt Parser. This table contains the percent identity, alignment length, and number of mismatches of each unique read ID against both the cymt and numt reference sequences. The output table also contains a classifier that tags each read as putatively originating from cymt or numt based on the higher percent identity or tagged as “undetermined” when reads have equal percentage identity to both the cymt and numt pseudogene references. Read_n indicates the *n*-th read in the dataset.

| read_ID<sup>1</sup> | read_in_pair<sup>2</sup> | mt_aln_bp<sup>3</sup> | mt_mismatch<sup>4</sup> | mt_identity<sup>5</sup> | numt_aln_bp<sup>6</sup> | numt_mismatch<sup>7</sup> | numt_identity<sup>8</sup> | candidate<sup>9</sup> |
| --- | --- | --- | --- | --- | --- | --- | --- | --- |
| Read_1 | 0 | 100 | 2 | 0.980000 | 97 | 5 | 0.948454 | cymt |
| Read_2 | 0 | 74 | 2 | 0.972973 | None | None | None | cymt |
| Read_3 | 0 | 100 | 2 | 0.980000 | 100 | 1 | 0.990000 | numt |
| Read_4 | 0 | 93 | 1 | 0.989247 | None | None | None | cymt |
| Read_5 | 0 | 100 | 0 | 1.0000 | None | None | None | cymt |
| Read_n | 0 | 100 | 0 | 1.0000 | 100 | 0 | 1.0000 | undetermined |

<sup>1</sup>Read identifier; <sup>2</sup>Orientation of reads in the pair (0 for single-end reads); <sup>3</sup>Length of alignment to cymt (bp); <sup>4</sup>Number of bp mismatches between read and cymt reference; <sup>5</sup>Percentage identity between read and cymt reference; <sup>6</sup>Length of alignment to numt (bp); <sup>7</sup>Number of bp mismatches between read and numt reference; <sup>8</sup>Percentage identity between read and numt reference; <sup>9</sup>Putative DNA from which the read originated.

### Post Numt Parser processing

Numt Parser produces an output table that lists per-read statistics for alignments to the cymt and numt reference (Table 1). This output table can be used to identify a subset of reads to retain for further analysis, allowing the user to subset the sequenced reads into a dataset most relevant for their study, and also allows for the compilation of datasets that include only reads originating from numts. For example, in this study we used the output table to compile a list of read IDs that were classified as “cymt” or “Undetermined” candidates. We then used Picard Toolkit v. 2.10.1 to select and retain only reads included in the read ID list and exported the output in BAM format. Coverage in the resulting datasets were calculated using SAMtools `depth`. Mitogenome alignment statistics (e.g., loss of coverage and mismatches across the genome pre- and post-filtering with Numt Parser) were averaged using a gaussian kernel-smoothing algorithm implemented in the `ksmooth` function in the statistical software R (R Core Team, 2019).

### Numt filtering using alternative bioinformatic approaches

In addition to the use of primers that prevent numt amplification (Curry et al., 2019), previous studies (Curry et al., 2021) have also used bioinformatic approaches that relied solely on read alignment properties to account for numt DNA contamination in lion datasets. We compared Numt Parser to this published approach, and to a second approach that used Basic Local Alignment Search Tool (BLAST) to categorize reads as putatively cymt or numt in origin.

#### Alternative bioinformatic approach 1: BWA and SAMtools

We filtered reads based on the alignment properties to a concatenated reference file that contained both cymt and numt reference sequence (hereafter referred to as SAMtools filtering). This approach has previously been used to account for numt contamination when reconstructing lion mitogenomes (Curry et al., 2021). Using the `mem` module of BWA, we aligned reads to a single FASTA file containing both cymt and numt sequences. Alignments were then processed with SAMtools `view` to remove unmapped reads, alignments with a mapping quality less than 30, secondary, and supplementary alignments. This filtered dataset was composed of only high-quality alignments that primarily mapped either to the cymt or numt references. The processed and indexed alignments were further filtered using SAM`TOOLS view` to exclude reads mapped to the numt reference, effectively keeping only reads of true cymt origin. Coverage for this filtered dataset was then calculated using the SAM`TOOLS depth` function.

#### Alternative bioinformatic approach 2: Basic Local Alignment Search Tool (BLAST)

Numt Parser was also compared against an approach that filters the dataset using BLAST (hereafter referred to as BLAST filtering). A local nucleotide BLAST database was constructed from a single reference FASTA file composed of both cymt and numt reference sequences using `makeblastdb` v2.4.0+ (Camacho et al., 2009). The reads of JCK 10711 and JCK 10712 were separately matched against this database using `blastn` v2.4.0+ and hits with an e-value of at least 0.001 and a maximum of 10 hits per query were saved in tabular format (`outfmt 6`). The resulting output table was then filtered to obtain the top BLAST hit per read using a custom Python script which compared e-values from all hits for a given read query (see Data Availability). If a top hit was the result of a comparison to the cymt reference sequence, the read query was returned as being putatively of cymt origin. From the resulting list of read IDs, the original alignment file for each sample was filtered using Picard Toolkit to generate a cymt-only BAM file. Coverage for this filtered dataset was then calculated using the SAM`TOOLS depth` function.

### Patterns of sequence disagreements

For each filtering method, mitochondrial consensus sequences were generated using the aligned reads from Cape lion samples JCK 10711 and JCK 10712. Variant sites (those differing from the reference cymt genome) were identified and filtered, using a minimum per-base quality of 20 and minimum coverage of 3X, in BCFtools (Danecek et al., 2021) and BEDtools (Quinlan & Hall, 2010). The coverage cut-off was implemented to increase accuracy of variant calls considering DNA damage and low DNA quality associated with ancient and historic samples (Parks & Lambert, 2015). The resulting consensus sequences for each sample were individually aligned to the cymt reference (KP202262) using the MAFFT aligner v7.310 (Katoh & Standley, 2013). For comparative purposes, we also aligned the consensus built from the numt-contaminated dataset against the lion cymt reference. The resulting alignments were used to visualize the sequence disagreements (i.e., differences due to a combination of biological polymorphism and technical error) between a given consensus sequence and the reference (Figure 2B; Figure 3). In addition, we aligned the unfiltered and Numt Parser-filtered consensus sequences for Cape lions JCK 10711 and JCK 10712 against each other in order to observe the retention of variant sites between the two individuals (Figure 2C).

**Figure 3.**
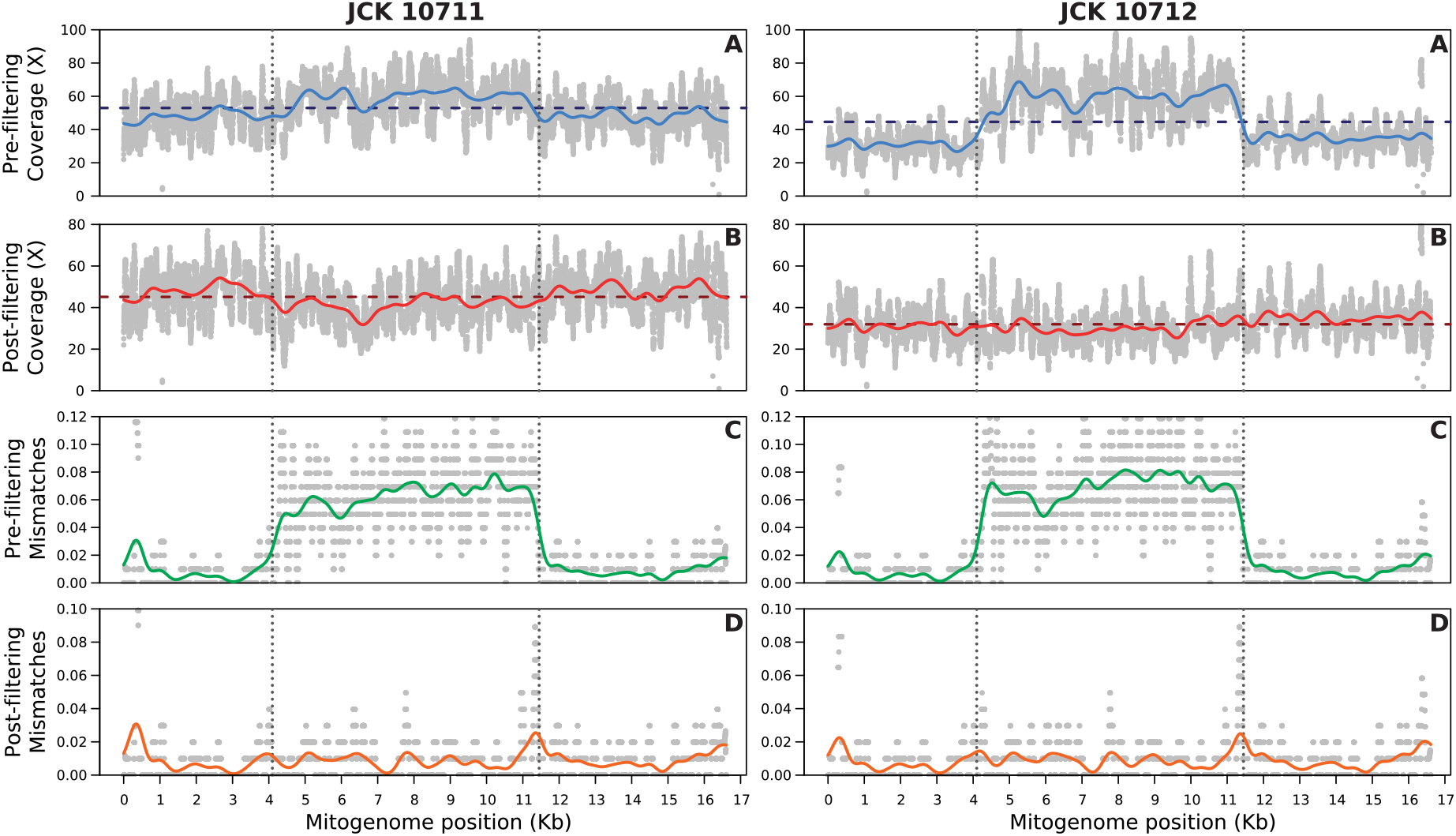
Patterns of read coverage and nucleotide mismatches spanning the two Cape lion mitogenomes, JCK 10711 (left panels) and JCK 10712 (right panels). The top row (A) shows the depth of coverage (X-fold reads) in the unfiltered dataset for JCK 10711 (left) and JCK 10712 (right). Grey dots show the per-base coverage, and the blue lines represent the kernel-smoothed coverage, averaged across a 500 bp sliding window. The horizontal dashed line shows the mean pre-filtering coverage for each dataset. Vertical dotted lines in all plots denote the start and end positions of the 7.2 kb region for which a numt pseudogene has been characterized in lion (Li et al., 2016). Row B shows the depth of coverage for each lion after filtering each dataset with Numt Parser. Grey dots show the per-base coverage, the red lines represent the kernel-smoothed average coverage across a 500bp sliding window, and the horizontal dashed lines show mean coverage values for each mitogenome. Row C shows the frequency of mismatches (i.e., nucleotide disagreements) in the alignment between the consensus sequence generated from the unfiltered reads and the lion cymt reference sequence. The grey dots show the frequency of mismatches per-100 bp. The green line shows the kernel-smoothed average frequency of mismatches per-100 bp, averaged across 500 bp sliding windows. Row D shows the frequency of nucleotide mismatches in the alignment between the consensus sequence derived from Numt Parser filtered datasets and the lion cymt reference. Grey dots show the frequency of mismatches per-100 bp, and the orange line shows the kernel-smoothed average frequency of mismatches per-100 bp. The unprocessed data for both JCK 10711 (left panel) and 10712 (right panel) shows an increase in coverage (A) and nucleotide disagreements (C) limited to the region spanning the characterized 7.2 kb lion numt. For both datasets, the distribution of coverage (B) and nucleotide mismatches (D) is equalized across the mitogenome as a consequence of filtering with Numt Parser.

## Results

The Cape lion DNA damage patterns showed nucleotide base-pair changes characteristic of ancient DNA, e.g., deamination of cytosine to uracil (Hofreiter, Jaenicke, et al., 2001). For both Cape lions, we observed a pattern of above average coverage that corresponded in location on the mitogenome with the 7.2 kb lion-specific numt pseudogene identified by Li et al. (2016). JCK 10712 exhibits a greater increase in coverage in the numt region (Figure 3A, right panel) than JCK 10711 (Figure 3A, left panel), which suggests that JCK 10712 has a higher degree of numt dataset contamination. Although the Cape lions have different patterns of above average coverage, both JCK 10711 and JCK 10712 (Figure 3C) exhibit a drastic increase in sequence mismatches in the region corresponding to the lion-specific numt pseudogene. Therefore, even for datasets that have comparatively low amounts of numt contamination based on above average coverage, e.g., JCK 10711 compared to JCK 10712, numt contamination may still impact the frequency of sequence mismatches in the consensus mitogenome sequence. Despite the differences observed in the two Cape lion datasets, the co-localization of both patterns of above-average coverage and elevated sequence disagreements within the span of a previously characterized lion-specific numt (Li et al., 2016) are strong indicators of the presence of numt contamination in these datasets.

### Numt Parser filters out reads of numtDNA origin without compromising biological signal

After filtering out reads of putative numt origin using Numt Parser, the loss of read coverage was limited to the 7,200 bp region that corresponds to the numt previously identified in lions (Figure 3B). We observed a greater loss of coverage across the numt region in JCK 10712 than in JCK 10711, consistent with having different levels of numt contamination (Table 2). Numt Parser therefore allows the user to equalize the average coverage across the genome by selectively removing reads of putative numt origin, without a substantial loss of read coverage from reads of cymt origin (Figure 3B).

**Table 2.** Alignment statistics for Cape lion mitochondrial genomes before and after filtering with Numt Parser.

| Sample | Before NUMT PARSER filtering |  |  | After NUMT PARSER filtering |  |  |
| --- | --- | --- | --- | --- | --- | --- |
|  | Reads (n) | Avg. cov (X) | Breadth of coverage (%) | Reads (n) | Avg. cov (X) | Breadth of coverage (%) |
| JCK 10711 | 13,729 | 52.99 | 100 | 11,879 | 45.13 | 100 |
| JCK 10712 | 10,891 | 44.63 | 100 | 8,070 | 31.98 | 100 |

When comparing the consensus sequences against the cymt reference (Figure 2B; Figure 3C), numt dataset contamination is evident as an overall increase in sequence disagreements (due to polymorphism and technical noise), as would be expected of a dataset with numt contamination (Li et al., 2016). When comparing our Cape lion sequences before and after processing with Numt Parser, we found that filtering led to a substantial reduction in the number of disagreements to the reference across both samples (Figure 2B, rows 3 and 7; Figure 3D). This reduction points to the removal of numt reads from the sequence pool, resulting in an uncontaminated mitogenome dataset.

Following Numt Parser filtering, the alignment between the filtered JCK 10711 and JCK 10712 consensus sequences still showed evidence of some disagreements (Figure 2C), indicating that informative nucleotide base differences were retained despite the filtering process. Moreover, the filtered mitogenome consensus sequences for JCK 10711 and JCK 10712 are different, highlighting that the two Cape lions had distinct mitochondrial haplotypes. Numt Parser therefore allows for the removal of disagreements due to numt contamination but concomitantly retains informative cymt nucleotide base differences, thus not sacrificing the true biological signal in the data.

### Numt Parser outperforms available filtering alternatives

For all three filtering approaches, including Numt Parser, SAMtools and BLAST filtering, the filtering of datasets to exclude reads of putative numt origin resulted in a greater loss of coverage for JCK 10712 (Figure 4B) than for JCK 10711(Figure 4A), consistent with JCK 10712 having greater numt contamination. However, post-filtering coverages differed between the three bioinformatic approaches. The mitochondrial-wide average coverage for JKC 10711 and JKC 10712, respectively, were 52.99x and 44.63x for the unfiltered datasets, 45.13x and 31.98x after Numt Parser filtering, 40.80x and 28.51x after SAMtools filtering, and 42.32x and 30.33x after BLAST filtering (see Supplementary Table 1). The same trend is evident when comparing the number of uncalled sites, i.e., sites with less than 3x coverage pre- and postfiltering. For JKC 10711 and JKC 10712, the unfiltered and Numt Parser filtered datasets each contained 1 and 2 uncalled sites, respectively. By contrast, SAMtools filtering resulted in 245 and 340 uncalled sites, while BLAST filtering resulted in 37 and 38 uncalled sites, respectively (green bars in Figure 2B; Supplementary Table 1).

**Figure 4.**
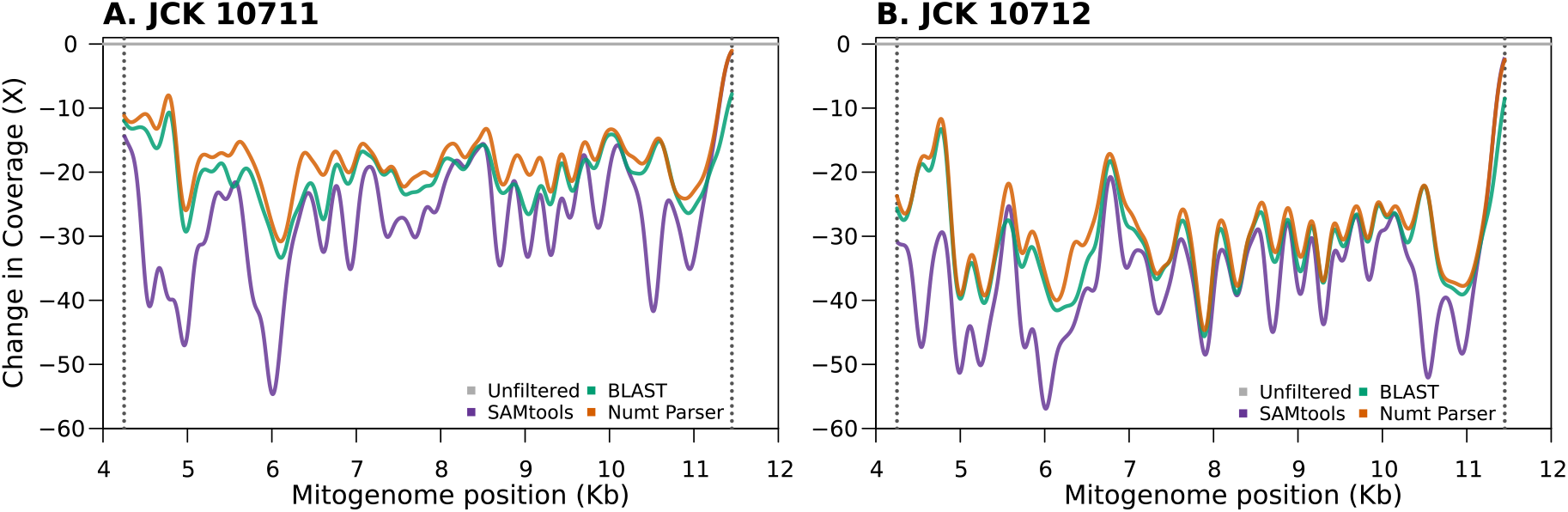
Three different bioinformatic filtering approaches (Numt Parser, SAMTools, BLAST) yielded consensus sequences with different coverages for JCK 10711 (A, left) and JCK 10712 (B, right). Coverage is only shown for positions within the defined 7.2 kb numt region (x-axis). The y-axis shows the reduction in coverage of the filtered dataset against the original, unfiltered dataset (horizontal grey line). The lines show the kernel-smoothed reduction in coverage, averaged across 250 bp sliding windows. The coverage loss is greater for the SAMtools filtered (purple) dataset than for BLAST filtered (green) and Numt Parser filtered (orange) datasets.

The greater loss of coverage for SAMtools and BLAST filtering than for Numt Parser filtering was especially evident in the region spanned by the lion numt pseudogene (Figure 4; Supplementary file 2). Comparing pre- and post-filtering coverages for the 7.2 kb numt region, the SAMtools filtered dataset (Figure 4, purple) had the largest decrease in coverage. This marked decrease was evident also for JKC 10711 (Figure 4A, purple) which had lower levels of numt contamination, for which SAMtools filtering resulted in a coverage reduction of 27.52x (Supplementary Table 2).

Within the numt region, loss in coverage between pre- and post-filter datasets for Numt Parser and BLAST was substantially less than for the SAMtools filtering approach (Figure 4). An average reduction of 32.16x was observed for SAMtools, versus reductions of 25.96x and 23.58x for BLAST and Numt Parser respectively (Supplementary Table 2). Numt Parser and BLAST filtering both resulted in consensus sequences that had fewer disagreements to the lion cymt reference compared to the unfiltered datasets while still retaining informative sites (Figure 2C); however, Numt Parser had the highest coverage post filtering for both JKC 10711 and JKC 10712.

## Discussion

Our software Numt Parser helps eliminate numt sequences in contaminated mitogenome datasets by identifying sequencing reads that likely originate from the numt pseudogene. These contaminating numt sequences can then be removed, resulting in filtered datasets that more accurately represent the true sequence of the mitogenome. Here we demonstrated the effectiveness of Numt Parser for removing numts when reconstructing the mitogenomes of two extinct Cape lions, and show that Numt Parser outperforms other bioinformatic filtering approaches. Numt Parser achieves these results by retaining more cymt reads to yield higher average genome coverages post-filtering, while minimizing numt dataset contamination. Retaining as many reads as possible without compromising the accuracy of the consensus sequence is especially pertinent for studies on ancient DNA where endogenous DNA copy numbers are often low and where low-depth genomic datasets are standard.

Numt Parser filtering resulted in less read coverage loss than SAMtools filtering (Figure 4). Due to this difference in loss of coverage, the number of uncalled sites in the consensus sequences were two orders of magnitude higher after SAMtools filtering than after Numt Parser filtering (Supplementary Table 1-2). SAMtools filtering likely discarded cymt-derived reads, causing the loss of informative variant sites. The SAMtools-based method processes alignments against a single FASTA containing both cymt and numt reference sequences. The presence of two simultaneous references on which to map the reads might obfuscate the identification of their true origin. Reads of cymt origin can still align to the numt reference with high mapping qualities (MAPQ), and *vice versa*, making the identification of primary alignments more difficult. When reads map with equal mapping quality to both references, cymt reads will be discarded when filtering out reads that have multiple alignments, reducing the overall coverage and the downstream generation of consensus sequences. The Numt Parser approach of independently mapping to cymt and numt sequences is therefore more effective in identifying reads as cymt or numt, and thus in retaining cymt reads.

Filtering efficacy also explains the success of Numt Parser over the BLAST-based approach. While the differences in coverage are not as substantial as for SAMtools (Figure 4), BLAST filtering did result in an order of magnitude increase in uncalled sites (Supplementary Table 1-2) when compared to Numt Parser filtering. Interestingly, the majority of reads lost when filtering with BLAST were not due to alignment filtering (i.e., reads discarded due to poor values of sequence similarly, alignment length, or e-value). Instead, it appears that a portion of reads are unable to be aligned by BLAST prior to filtering. By default, the BLAST+ suite uses the `megablast` algorithm when performing nucleotide-to-nucleotide alignments (Camacho et al., 2009; Madden, 2020), which requires an exact match of at least 28 bp to align two sequences (controlled by the -`word_size` parameter). This alignment requirement is too stringent for short reads – for which BLAST has never been the primary alignment tool – particularly those originating from aDNA. For example, when aligning with `blastn` using default settings, we observe that only 93.86% and 95.66% of reads are aligned for JKC 10711 and JKC 10712, respectively. By decreasing the `word_size` parameter to 11, matching that of -`task blastn`, the proportion of alignments can be further increased to 99.54% for JKC 10711 and 99.59% for JKC 10712. However, this comes at the cost of obtaining more small, spurious alignments. Therefore, while we found that increasing the performance of the BLAST filtering is possible, it does require the use of non-default parameters. This behavior, in addition to the external script required for parsing and filtering the resulting alignments, further limits the BLAST filtering approach when compared to Numt Parser.

Numt Parser relies on well-defined numt and cymt reference sequences (such as those available for the lion) for both alignment and sequence identity comparisons. It additionally assumes that the cymt reference is authentic and was not reconstructed using a numt contaminated dataset. These requirements also apply to any alignment-based method for numt filtering, including both SAMtools and BLAST filtering methods presented in this paper. The accurate characterization of numts can be cumbersome for non-model organisms. Numt pseudogene reconstruction and characterization may be approached through molecular laboratory and bioinformatic methods (Bensasson et al., 2001; Smart et al., 2019). For example, DNA extractions can be enriched for nDNA instead of cymt before PCR amplification, e.g., by using nDNA rich tissues (Zhang & Hewitt, 1996; Zischler, Geisert, et al., 1995). Alternatively, genomic library construction and sequencing technologies that rely on single molecule, long read sequencing (e.g., Nanopore and PacBio) may be used to identify cymt mitogenomes and concomitantly also numt pseudogenes (Vossen & Buermans, 2017). For numt pseudogenes in particular, longer reads may be especially beneficial, as they can be used to identify integration sites in the nuclear genome, improving both the differentiation between numt and cymt sequences, and the inclusion of numt pseudogenes in nuclear reference genomes (Sohn & Nam, 2018). Bioinformatic approaches have also (Chakraborty et al., 2019) been developed to identify and characterize numts, e.g., by comparing differences in the topological properties of haplotype networks based on cymt and numt sequences (Smart et al., 2019).

While our implementation of Numt Parser focused on identifying and removing numt contamination in *Panthera*, the opposite application is also possible. Comparing the sequencing reads against a characterized numt reference can instead be used to identify, retain, and compile a dataset of numt reads. Numt Parser may therefore enable the use of numt pseudogenes as genetic markers for evolutionary inferences. For example, numt pseudogene sequences may be important genetic markers for the discovery and characterization of ancient mitochondrial haplotypes (Lammers et al., 2017). Numt pseudogenes have been applied to reconstruct ancestral mitochondrial sequences in the polar bear (*Ursus maritimus*), where historical mitogenome sequences are thought to be lost in extant populations, but putatively preserved as numt pseudogene paralogs (Lammers et al., 2017). Numt pseudogene sequences may also be informative in taxa where they represent DNA transfer events that predate speciation. In such taxa they may provide a model of evolutionary change that could potentially be used to resolve phylogenetic inconsistencies, e.g., in taxa where incomplete lineage sorting or hybridization occurred (Wang et al., 2015). The ability to better characterize numt pseudogene sequences may also benefit biomedical applications. For example, distinguishing numt pseudogene copies from true cymt mitogenomes can help minimize inaccurate characterization of pathogenic mitochondrial variants and can contribute towards our understanding of mitochondrial diseases (Calvignac et al., 2011; Hazkani-Covo et al., 2010; Wallace et al., 1997).

We tested Numt Parser using shotgun sequenced data from two ancient Cape lions, showing that Numt Parser is effective at removing numt contamination in lions. Numt Parser should be effective at removing numt contamination for other species within the genus *Panthera*, which possess well characterized numts (Li et al., 2016), and for other taxa for which numt pseudogene reference sequences are available. In lions and other *Panthera*, there are a large number of long tandem numt repeats known to have interfered with accurate mitogenome reconstruction (Li et al., 2016). Numt Parser may be especially helpful in removing contamination by such tandem numt repeats, as high copy number may increase numt-associated read coverage and consequently also sequence disagreements.

In summary, Numt Parser allows for the identification and removal of numt contamination in high-throughput sequencing data, enabling mitochondrial genome reconstruction. Numt Parser equalizes coverage across the mitogenome by removing reads that originate from numts, decreasing the patterns of sequence disagreements that arise from numt contamination. The assessment and removal of reads originating from numts through the use of Numt Parser can improve the reconstruction of mitogenomes, allowing for more accurate and robust biological inferences.

## Acknowledgements

For funding, we thank the USAID Wildlife TRAPS Project and the UIUC ACES Office of International Programs. AdeF was supported by the Program in Ecology, Evolution and Conservation Biology Research Award, UIUC, and by the Cooperative State Research, Education, and Extension Service, US Department of Agriculture, under project number ILLU 875–952. AGR-C was supported by NSF grant 1645087. We thank the UIUC High-Throughput Sequencing and Genotyping Unit. We thank Dr. Tolulope Perrin-Stowe for her help in collecting the ancient DNA samples at the FMNH. We thank Hazel Singer and the rest of the family of the late Dr. R Singer (University of Chicago) for access to and use of these historical specimens.

## Data availability

The Cape lion mitogenome sequences are available on GenBank under accession numbers XXXXXXXX-XXXXXXXX.

Numt Parser GitHub repository: https://github.com/adeflamingh/NuMt_parser

