## Supplemental file 1 for "NUMT PARSER: automated identification and removal of nuclear mitochondrial pseudogenes (numts) for accurate mitochondrial genome reconstruction in *Panthera*"

### de Flamingh et al - aDNA extraction and genomic library protocol

The information below describes the protocol for extracting whole genome DNA from ancient or degraded samples, and building genomic libraries for Illumina sequencing.

### Samples

1. JCK 10711 - sample A
2. JCK 10711 - sample B
3. JCK 10712 - sample A
4. JCK 10712 - sample B
5. Extraction Negative
6. Library Negative

#### Reagents and Equipment:

##### Day 1: sample drilling and digestion

- Dental burs Weigh boats Isopropanol Bleach DNA-Off
- 15 mL polypropelene tubes
- 0.5M EDTA
- 3.33mg/mL Proteinase K
- 10% N-lauryl sarcosine

##### Day 2: DNA extraction

- 15mL centrifugal filter units (30K molecular weight filter)
- Qiaquick PCR Purification Kit:
  - Qiaquick minicolumn/collection tube PB Buffer
  - PE Buffer
  - Elution Buffer
- one set of standard 2mL tubes and one set of Qiagen 1.5mL tubes and one set of 1.5mL safe lock tubes
-

##### Day 3: genomic library construction

- [NEB II Ultra Illumina library kit with purification beads]
- [NEB Unique Dual Indexing primers for Illumina]
- Lo-Bind Eppendorf Tubes
- 15 mL polypropelene tube for the 80% ethanol wash
- 0.2ml PCR tubes (1 per sample)

#### Protocol

##### DAY 1:

- All equipment (including Dremel tool/drill), pipettors and tubes, must be decontaminated under UV light for a minimum of 15 minutes prior to use
- Change pipette tips after each use

STEP 1: CLEANING

1. Option 1 (sample collection in aDNA laboratory):
   a. Label weigh boats with sample names
   b. Soak up to 7 teeth or bone fragments in 40% bleach for 3 minute
   c. Rinse three times with diH20

d. Rinse once with isopropanol
e. Dry samples under UV in DNA Crosslinker for 25min or until completely dry

2. Option 2 (sample at museum/off-site):
   a. Decontaminate work space, Personal Protective Equipment (PPE) and other equipment with Takara Bio DNA-off

   - PPE: gloves, disposable gown/labcoat, sleeves and mask
   - Equipment should include UV decontaminated drill burs and all other items needed to collect sample (e.g. weigh boats; 15 mL polypropelene tubes )
     b. Place sample in decontaminated weigh boat, wipe down the area on the bone/tooth where sample will be collected with 6% sodium hypochlorite (full strength Clorox bleach)
     c. Remove all dirt from tooth or bone surface
     d. Rinse three times with diH20
     e. Dry area with sterile paper towel (e.g. Kimwipe) and leave to further airdry for 5 minutes.

STEP 2: DRILLING

1. Clean the inside of the drilling hood using bleach or DNA-Off (if sampling in aDNA laboratory)

2. Attach a UV decontaminated dental bur to the Dremel tool/drill

3. Place sheet of UV decontaminated aluminum foil inside the drill hood (this allows for the easy collection of bone/tooth powder)

4. Drill a hole in the root of the tooth (or bone) until around 0.20g of sample is obtained

5. If drilling is ineffective, break off the end of the root of the tooth and crush using the mortar and pestle (bleach and UV-decontaminate before and after use)

6. Place powdered sample in 15ml polypropelene tubes and record the weight. Save excess sample in a separate decontaminated 15ml tube.

---

**SAMPLE PROCESSING FROM HERE ONWARDS MUST OCCUR IN AN ANCIENT DNA LABORATORY/FACILITY**

---

STEP 3: DIGESTION

1. Prepare 1 additional tube without biological sample (use molecular grade H2O instead) as the extraction negative control
2. Add 4ml of 0.5M EDTA to the 15ml tubes with powdered sample
3. Add 100ul of 33.3mg/ml proteinase K
4. Add 300ul of 10% N-lauryl sarcosine
5. Seal the lids of the tubes with parafilm
6. Place 15ml tubes in the rotating incubator, turn on the rotor and incubate 20-24hrs at 37°C or until all sample is digested

---

**DAY 2:**

STEP 4: CENTRIFUGAL CONCENTRATION

1. Label centrifugal filters with sample names
2. Spin 15 mL tubes down at 3500 rpm for 5 minutes to concentrate bone powder at the bottom
3. Add the digested sample supernatant to the centrifugal filter (try to avoid getting bone/tooth fragments in sample to speed centrifugation)
4. Spin at 3000-4000 rpm until concentrated to 250ul (approximately 35-50mins)
5. Pipette concentrated sample into a new 2ml tube (UV decontaminated)

STEP 5: DNA EXTRACTION PROTOCOL

*protocol uses Qiaquick PCR Purification kit, however, steps included in the original protocol have been modified*

1. Preheat incubator to 37'C

2. Label one set of standard 2mL tubes and one set of Qiagen 1.5mL tubes and one set of 1.5mL safe lock tubes with sample names (all tubes should be UV decontaminated prior to use)

3. Add 5 volumes of PB buffer (1250 uL or 2X625ul) to the amount of concentrated sample and vortex for 30s (pH indicator not necessary)

4. Label Qiaquick minicolumn and collection tube and add 740ul PB buffer mix to center of Qiaquick membrane

5. Incubate for 5 minutes at room temperature (not on heat block)

6. Spin at 13,000 rpm for 1 min and discard the flow through (flow through contains PB buffer and must be discarded appropriately, check with your facility/institution)

7. Repeat until all PB buffer mix has been filtered

8. Add 740ul PE buffer* to the center of the minicolumn

    - make PE buffer fresh using 1ulPE to 5ul 100% ethanol; make double the amount because you will need to complete this step twice Example: For 4 samples,4x2x750=6000 (1000ul PE & 5000ul ethanol)

9. Spin at 13,000 rpm for 1 min

10. Discard solution and centrifuge for an additional minute

11. Discard collection tube and place minicolumn into a clean, labeled 1.5ml

12. Add 30ul Elution buffer to the center of the QIAquick membrane

13. Incubate at 37°C in the heat block for 5min (cover with foil to ensure stable incubation temperature)

14. Centrifuge at 13,000rpm for 1 min

15. Add 30ul Elution buffer to the center of the QIAquick membrane

16. Incubate at 37°C in the heat block for 5min (cover with foil to ensure stable incubation temperature)

17. Centrifuge at 13,000rpm for 1 min

18. Discard the minicolumn and save the 1.5ml tube containing extracted DNA

STEP 6: REPEAT EXTRACTION TO REMOVE INHIBITORS

- Repeat from 5.3 to 5.18. For step 5.3, the starting volume is 30 +30 elution buffer, so the amount of PB buffer to add is 300uL.
- During this round of DNA extraction please use pipette to mix samples rather than vortexing

---

**DAY 3:**

**GENOMIC LIBRARY CONSTRUCTION PROTOCOL**

- All equiptment, including pipettors and tubes, must be decontaminated under UV light for a minimum of 15 minutes prior to use
- Change pipette tips after each use (do not suck up liquid with the same pipette tip more than once)
- Starting Material: 5 ng–1 µg fragmented DNA.
- use only [DNA LoBind tubes] to maximize nucleic acid recovery

**For genomic library construction with USER (Uracil-Specific Excision Reagent) pre-treatment follow Step #7A. For genomic library construction without USER pre-treatment follow Step #7B.**

STEP 7A: GENOMIC LIBRARY CONSTRUCTION WITH USER PRE-TREATMENT

a) Mix the following components in a sterile nuclease-free LoBind tube:

- 6.5 ul End Repair Reaction Buffer (10X)
- 55.2 µl Fragmented DNA (from Step #6)
- ·0.3 µl USER enzyme

b) Mix by pipetting followed by a quick spin
c) Incubate at 37'C for 3 hours (foil over incubator)
d) Add 3.0 ul EndPrep Enzyme Mix to each tube. Total volume per tube should be 65ul.
f) Mix bypipetting followed by a quick spin to collect all liquid from the sides of thetube.
Continue to Step #8

STEP 7A: GENOMIC LIBRARY CONSTRUCTION **WITHOUT** USER PRE-TREATMENT

a) Mix the following components in a sterile LoBind nuclease-free tube:

- 3.0 µl EndPrep Enzyme Mix
- 6.5 µl EndRepair Reaction Buffer (10X)
- 55.5 µl Fragmented DNA (from Step #6)

---------Total volume 65 µl

b) Mix bypipetting followed by a quick spin to collect all liquid from the sides of thetube.
Continue to Step #8

STEP 8: INCUBATION

a) Place sample in a heat block/incubator, with Aluminum Foil as a cover, and run the following program:
30 minutes at 20°C(Room Temp – not on heatblock)
30 minutes at 65°C(On heatblock; cover with foil)

b) Make a 1:25 Adaptor Dilution of adapter:molecular grade water to use in STEP 9

STEP 9: ADAPTER LIGATION

*Vortex and spin down all reagents before use*

a) Add the following components directly to the End Prep reaction mixture (65 µl )and mix well:

Blunt/TA Ligase Master Mix 15µl
NEBNext Adaptor for Illumina 2.5µl
Ligation Enhancer 1µl

------------------ Total volume 83.5 µl

b) Mix by pipetting followed by a quick spin to collect all liquid from the sidesof the tube.
c) Incubate at 20°C for 15 minutes (Room Temp)
d) Add 3 µl of USER enzyme to the ligation mixture
e) Mix well and incubate at 37°C for 15minutes.

STEP 10: CLEANUP OF ADAPTER-LIGATED DNA WITHOUT SIZE SELECTION

- Follow standard steps described in the NEB protocol for cleanup of adapter-ligated DNA without size selection (for input < 50ng DNA)

STEP 11: SETUP PCR AMPLIFICATION OF ADAPTER-LIGATED DNA

a) setup a PCR reaction by mixing the following components in sterilestrip tubes:

Adaptor Ligated DNA Fragments 15 µl
NEBNext High Fidelity Master Mix 25 µl
Unique Dual Index Primer Pairs 10 µl
-------------------- Total volume 50 µl

---

#### Samples are transported to a different facility for PCR and further processing

---

---

STEP 12: PCR (ROUND 1)

| CYCLE | TEMPERATURE | TIME | #CYCLES |
|---|---|---|---|
| Initial denaturation | 98'C | 30sec | X1 |
| – | – | – | – |
| Denaturation | 98'C | 10sec | 4-12 (12 maximum) |
| Annealing/Extension | 65'C | 75sec | |
| – | – | – | – |
| Final Extension | 65'C | 5minutes | X1 |
| – | – | – | – |
| Hold | 4'C | | Infinity |

STEP 13: PCR (ROUND 2)
a) Set up four PCR reactions for each sample using PCR (round 1) as template DNA and the following components

*Vortex reagents before use, quick spin down *
*Use high quality Taq polymerase (e.g. Phusion HF Taq)*

Master Mix (e.g. Phusion HF Taq) - 25µl
H2O - 14.5µl
IS5 Primer (Meyers et al 2010) - 1.5µl
IS6 Primer - 1.5µl
DMSO - 1.5µl
BSA - 1µl
DNA (from PCR1) - 5µl
-------Total volume 50

Freeze the remaining PCR product from round1 at -20'C.
** PCR reactions are split to increase read diversity of amplified fragments and decrease PCR duplication

b) Run the following PCR:

| CYCLE | TEMPERATURE | TIME | #CYCLES |
|---|---|---|---|
| Initial denaturation | 95’C | 4minutes | X1 |
| – | – | – | – |
| Denaturation | 95’C | 15sec | 12 cycles |
| Annealing/Extension | 65’C | 30sec | |
| Annealing/Extension | 68’C | 30sec | |
| – | – | – | – |
| Hold | 10’C | | Infinity |

STEP 14: MinElute CLEANUP OF AMPLIFIED PCR PRODUCT

- (Qiagen MinElute PCR Purification Kit)

1. Pool the 4 reactions for each library and add 5X the volume of PB Buffer (e.g. 4*50 uL = 200 uL so add 1000 uL of thePB Buffer).
2. Transfer 625 uL of the DNA/PB mix to each MinElute spin column
   Centrifuge for 1 minute at 13, 000 rpm. Remove the spin column from the collection tube, discard the flow-through and put the column back in the collection tube.
3. Repeat Steps 2 and 3 until all of theDNA/PB buffer mix has been filtered through.
4. Add 750 uL PE Buffer to the spincolumn. Centrifuge for 1 minute at 13, 000 rpm.
5. Empty the collection tube into the waste container, put the spin column back in the collection tube and return to centrifuge. Spin 1 additional minute at 13, 000rpm to dry the spin column filter
6. Discard the collection tube and place the spin column in a new, labeled 1.5 mL SafeLock tube.
7. Add 31.5 uL EB and incubate at 37’C for 5 minutes

** to maximize eluted DNA concentration repeat this step twice by adding 16 EB, incubating for 5 minutes, centrifuging for 1 min at 13, 000 rpm and repeating

8. Centrifuge for 1 minute at 13, 000 rpm.

9. Genomic libraries are now ready for fragment analysis and sequencing.
