## Supplemental file 2 for "NUMT PARSER: automated identification and removal of nuclear mitochondrial pseudogenes (numts) for accurate mitochondrial genome reconstruction in *Panthera*"

**Supplementary Table 1.** Sample names and complete mitochondrial genome coverage statistics for unfiltered, SAMTOOLS filtered, BLAST filtered, and NUMT PARSER filtered datasets. For each method, we report the number of nucleotide base-pairs (num.sites), the average depth of coverage as X-fold (avg.cov), the average change in consensus sequence coverage for different filtering approaches compared to unfiltered datasets (avg. change), the median and standard deviation (median.cov and stdv.cov), the minimum and maximum X-fold read coverage at each position in the mitogenome (min.cov and max.cov) and the number of sites in the genome that have less than 3 X-fold read coverage (low.cov.sites).

| sample.id | method | num. sites | avg. cov | avg. change | median. cov | stdv. cov | min. cov | max. cov | low.cov. sites |
|---|---|---|---|---|---|---|---|---|---|
| JKC 10711 | unfiltered | 16,620 | 52.99 | 0.00 | 53.0 | 12.43 | 1.0 | 94.0 | 1.0 |
| JKC 10711 | samtools | 16,620 | 40.80 | -12.19 | 43.0 | 14.91 | 0.0 | 78.0 | 245.0 |
| JKC 10711 | blast | 16,620 | 42.32 | -10.67 | 42.0 | 10.92 | 0.0 | 76.0 | 37.0 |
| JKC 10711 | numt_parser | 16,620 | 45.13 | -7.86 | 45.0 | 10.67 | 1.0 | 78.0 | 1.0 |
| JKC 10712 | unfiltered | 16,620 | 44.63 | 0.00 | 40.0 | 17.16 | 2.0 | 101.0 | 2.0 |
| JKC 10712 | samtools | 16,620 | 28.51 | -16.12 | 29.0 | 10.82 | 0.0 | 80.0 | 340.0 |
| JKC 10712 | blast | 16,620 | 30.33 | -14.29 | 30.0 | 8.73 | 0.0 | 75.0 | 38.0 |
| JKC 10712 | numt_parser | 16,620 | 31.98 | -12.65 | 31.0 | 8.76 | 2.0 | 82.0 | 2.0 |
| Average* | unfiltered | 16,620 | 48.81 | 0.00 | 46.5 | 14.80 | 1.5 | 97.5 | 1.5 |
| Average* | samtools | 16,620 | 34.65 | -14.16 | 36.0 | 12.87 | 0.0 | 79.0 | 292.5 |
| Average* | blast | 16,620 | 36.33 | -12.48 | 36.0 | 9.82 | 0.0 | 75.5 | 37.5 |
| Average* | numt_parser | 16,620 | 38.56 | -10.25 | 38.0 | 9.72 | 1.5 | 80.0 | 1.5 |

*for JKC 10711 and JKC 10712 combined

**Supplementary Table 2.** Sample names and numt pseudogene coverage statistics for unfiltered, SAMTOOLS filtered, BLAST filtered, and NUMT PARSER filtered datasets. For each method, we report the number of nucleotide base-pairs (num.sites), the average depth of coverage as X-fold (avg.cov), the average change in consensus sequence coverage for different filtering approaches compared to unfiltered datasets (avg. change), the median and standard deviation (median.cov and stdv.cov), and the minimum and maximum X-fold read coverage at each position in the mitogenome (min.cov and max.cov).

| sample.id | method | num.sites | avg.cov | avg.change | median.cov | stdv.cov | min.cov | max.cov |
|---|---|---|---|---|---|---|---|---|
| JKC 10711 | unfiltered | 7,201 | 59.375 | 0.000 | 60.0 | 12.0 | 23.0 | 94.0 |
| JKC 10711 | samtools | 7,201 | 31.851 | -27.525 | 33.0 | 14.6 | 0.0 | 69.0 |
| JKC 10711 | blast | 7,201 | 38.897 | -20.479 | 39.0 | 9.9 | 10.0 | 67.0 |
| JKC 10711 | numt_parser | 7,201 | 41.374 | -18.001 | 41.0 | 9.7 | 12.0 | 69.0 |
| JKC 10712 | unfiltered | 7,201 | 59.760 | 0.000 | 60.0 | 13.5 | 21.0 | 101.0 |
| JKC 10712 | samtools | 7,201 | 22.958 | -36.803 | 23.0 | 10.6 | 0.0 | 62.0 |
| JKC 10712 | blast | 7,201 | 28.860 | -30.901 | 28.0 | 8.6 | 9.0 | 67.0 |
| JKC 10712 | numt_parser | 7,201 | 30.609 | -29.152 | 29.0 | 8.9 | 10.0 | 67.0 |
| Average* | unfiltered | 7,201 | 59.568 | 0.000 | 60.0 | 12.7 | 22.0 | 97.5 |
| Average* | samtools | 7,201 | 27.404 | -32.164 | 28.0 | 12.6 | 0.0 | 65.5 |
| Average* | blast | 7,201 | 33.878 | -25.690 | 33.5 | 9.2 | 9.5 | 67.0 |
| Average* | numt_parser | 7,201 | 35.991 | -23.576 | 35.0 | 9.3 | 11.0 | 68.0 |

*for JKC 10711 and JKC 10712 combined

**Supplementary Figure 1.** Amino acid sequence alignments of genes within the 7.2 kp span of the previously characterized lion numt pseudogene. Top panel shows alignment of the coding sequence of the gene *ND2* between the lion cymt reference sequence (KP202262), the lion numt reference (KF907306), and the consensus sequences generated from the two Cape lion samples, JCK 10711 and JCK 10712 after NUMT PARSER filtering. Homologous sites with conserved amino acid residues are shown in dark gray. Sites with variable residues are shown in light gray, with sites highlighted in white being different from the consensus. Middle and bottom panels show alignments for the amino acid sequences of *COX1* and *COX3*, following the format described for the top panel. The amino acid positions for each gene are indicated as numbers above the coding sequences. For all three genes, the alignments show the presence of non-synonymous coding mutations present in the numt reference, evidence of pseudogenization. After filtering for numt contamination, the sequence for both Cape Lions shows sequences conserved to that of the lion cymt reference.

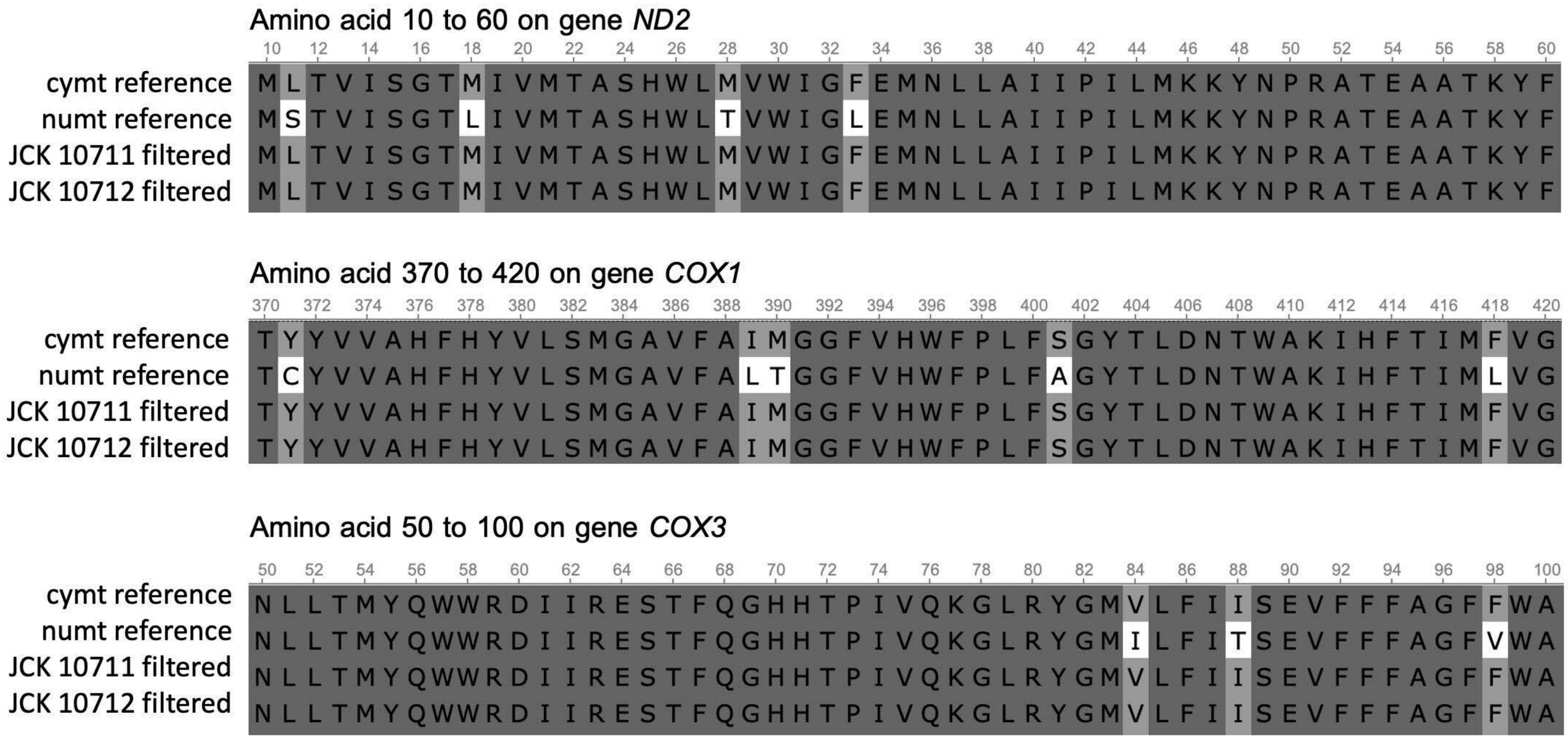
